# Genome-wide association study in a maize diversity panel dissects the genetic basis of lignocellulosic recalcitrance

**DOI:** 10.64898/2026.10.11.758201

**Authors:** Shaogan Wang, Frank Roeber, Susanne Groh, Therese Bolduan, Benjamin Stich, Delphine Van Inghelandt, Vicente Ramírez, Markus Pauly

## Abstract

Lignocellulosic biomass is an abundant, renewable resource for the production of biorenewable products. Its efficient utilization is largely hindered by the inherent recalcitrance of lignocellulose to degradation. Natural genetic variation in biomass saccharification efficiency and lignocellulose components across diverse maize germplasm provides a powerful resource to dissect the genetic basis of lignocellulosic recalcitrance. Here, we performed high-throughput profiling of lignocellulosic components and saccharification efficiency in mature stalks across a diverse panel of 239 field grown maize inbred lines, and identified extensive natural variation in biomass properties. Saccharification efficiency was strongly positively associated with matrix glucose content, moderately positively associated with the arabinose-to-xylose (Ara:Xyl) molar ratio and crystalline cellulose content, whereas saccharification efficiency was strongly negatively associated with xylose and lignin content. Genome-wide association studies (GWAS) were conducted to identify single nucleotide polymorphisms (SNPs) associated with saccharification efficiency and lignocellulosic components. Numerous significant single nucleotide polymorphisms (SNPs) were identified, defining 79 quantitative trait loci (QTLs) across all 10 maize chromosomes. The presence of pleiotropic SNPs and the extensive co-localization of QTLs for multiple traits highlight the complex genetic architecture and shared genetic basis of maize biomass properties. Our study reveals substantial natural variation in maize biomass digestibility and identifies multiple novel genetic loci underlying variation in saccharification efficiency and lignocellulose components.

## Introduction

Maize (*Zea mays* L.) is one of the most widely cultivated cereal crops worldwide. Beyond its primary role in producing grain for food, animal feed, and industrial derivatives, maize also generates substantial amounts of lignocellulosic biomass (Erenstein et al., 2022; Zabed et al., 2023). Maize biomass represents an abundant and renewable feedstock for the bioeconomy, such as the production of cellulosic biofuel, providing a sustainable alternative to fossil fuels (Aghaei et al., 2022; Kaur et al., 2025). As global demand for biorenewable products continues to increase (Boysen-Urban et al., 2026), improving the digestibility and bioconversion efficiency of maize biomass has emerged as an important objective in maize genetic improvement and breeding (Vanhevel et al., 2024).

For lignocellulosic biomass conversion, the structural polysaccharides need to be hydrolyzed into fermentable sugars through enzymatic saccharification (Houfani et al., 2020). However, the extensive covalent and non-covalent interactions among lignocellulosic polymers (i.e., cellulose, hemicellulose, and lignin) form a highly organized composite structure that provides mechanical plant strength and protection against pathogens while simultaneously limiting the accessibility of hydrolytic enzymes to the structural polymers (Zeng et al., 2017; Petridis and Smith, 2018). Consequently, biomass recalcitrance, defined as the intrinsic resistance of lignocellulosic biomass to enzymatic deconstruction, represents a major bottleneck for efficient biomass utilization (Himmel et al., 2007; Pauly and Keegstra, 2008).

Biomass recalcitrance is largely determined by the composition, architecture, and physicochemical properties of lignocellulose (Pauly and Keegstra, 2010; Zoghlami and Paës, 2019; Wang et al., 2021; Kirui et al., 2022). Cellulose microfibrils provide the principal mechanical framework, and their crystallinity, degree of polymerization, and spatial organization influence both wall strength and susceptibility to enzymatic hydrolysis (Arantes and Saddler, 2010; Meng and Ragauskas, 2014; Wu et al., 2021). Thus, genetic perturbation of cellulose biosynthesis and deposition alters wall mechanical properties while simultaneously affecting biomass digestibility (Li et al., 2017; Wu et al., 2019; Nayeri et al., 2022). However, alterations in cellulose content or structure often compromise plant growth and mechanical stability leading to lodging and reduced biomass yield (Zhang et al., 2021). Therefore, a pivotal challenge in current research is to genetically manipulate cellulose abundance to enhance saccharification efficiency without incurring detrimental effects on plant development, growth and thus yield (Sumiyoshi et al., 2013; Wang et al., 2025). Hemicelluloses are a heterogeneous group of polysaccharides characterized by β-(1→4)-linked glucan, xylan, or mannan backbones with diverse side-chain substitutions (Scheller and Ulvskov, 2010; Pauly et al., 2013). Through hydrogen bonding and other non-covalent interactions, hemicelluloses associate closely with the surface of cellulose microfibrils, forming a matrix that connects cellulose with other wall components and further constrains the accessibility of hydrolytic enzymes to cellulose (Jarvis, 2022; Østby and Várnai, 2023). Unlike dicotyledonous plants, the xylan backbones of grasses such as maize are extensively substituted with arabinose, glucuronic acid, and acetyl groups, forming glucuronoarabinoxylans (GAXs), the predominant hemicelluloses of grass walls (Smith et al., 2017a; Tryfona et al., 2023). The arabinosyl side chains on GAXs can be esterified with hydroxycinnamic acids, particularly ferulic acid (FA) and *p*-coumaric acid (*p*CA), enabling oxidative cross-linking between adjacent GAX molecules and between GAX and lignin, which generate a highly interconnected three-dimensional lignin-carbohydrate complex (LCC) network that reinforces the lignocellulose recalcitrance (Hatfield et al., 2017; Eugene et al., 2020; Xiao et al., 2025). Lignin constitutes a major barrier to enzymatic deconstruction. As a hydrophobic phenylpropanoid polymer, it restricts hydrolytic enzyme accessibility through physical encapsulation but also irreversibly binds cellulases via chemical adsorption (Gao et al., 2014). Genetic studies of classical *brown midrib* (*bm*) mutants and other maize mutants affected in their lignin biosynthesis have demonstrated that reducing lignin content or altering lignin composition can substantially enhance biomass digestibility and saccharification (Barrière et al., 2004; Smith et al., 2017b; Xiong et al., 2020; Vanhevel et al., 2024). Therefore, manipulating lignin biosynthesis is an important strategy to reduce wall recalcitrance (Halpin, 2019; Gui et al., 2020).

Emerging from the complex interplay of composition, organization, and physicochemical properties of multiple lignocellulosic components, biomass recalcitrance is inherently a quantitative trait controlled by numerous genetic loci (Lorenzana et al., 2010; Courtial et al., 2014; Torres et al., 2015; van der Weijde et al., 2017). In maize, forward genetic studies have identified a number of genes affecting recalcitrance (Xiong et al., 2019; Kraemer et al., 2021; Liu et al., 2021; Wang et al., 2023; Wang et al., 2024). However, these approaches typically focus on individual genes or mutations with relatively large phenotypic effects, thereby capturing a limited subset of the genetic loci underlying this complex trait. In contrast, genome-wide association studies exploit historical recombination events across natural populations, enabling high-resolution identification of genetic loci and candidate genes underlying these complex traits (Xiao et al., 2017; Shikha et al., 2021; Sahito et al., 2024). Several GWAS studies utilizing diverse maize germplasm panels of varying sizes have identified loci associated with variation in lignocellulosic components or saccharification efficiency, demonstrating the potential of leveraging natural variation to uncover genetic basis of maize biomass recalcitrance (Li et al., 2016; López-Malvar et al., 2019; López-Malvar et al., 2021; Gesteiro et al., 2025a; Gesteiro et al., 2025b; Ren et al., 2026). Despite these advances, few studies have systematically integrated comprehensive lignocellulosic compositional profiling with enzymatic saccharification to elucidate the genetic basis of biomass recalcitrance at maize natural population level.

Here, we combined high-throughput profiling of lignocellulosic components with enzymatic saccharification efficiency in mature maize stalks across a panel of 239 multi-location field grown maize inbred lines and performed genome-wide association analyses (GWAS) to identify genetic loci and candidate genes underlying natural variation in lignocellulosic components and biomass digestibility. Our results provide new insights into the genetic architecture of maize biomass recalcitrance and identify genetic resources that may facilitate the improvement of maize biomass utilization.

## Results

### Broad natural variation in cell wall digestibility and lignocellulosic components within a maize diversity panel

To evaluate the extent of natural variation in lignocellulosic-related traits within the maize germplasm, a panel of 239 inbred lines comprising Stiff Stalk (SS), Non-Stiff Stalk (NSS), and tropical germplasm was systematically evaluated for enzymatic digestibility (saccharification) alongside the absolute and relative content of major lignocellulosic components. Mature stalks were harvested from three field environments, processed to obtain de-starched alcohol-insoluble residue (dAIR), and subsequently subjected to enzymatic or acidic hydrolysis.

Lignocellulosic digestibility was quantified via enzymatic saccharification efficiency (Sac), determined by measuring the glucose yield (mg L⁻¹) released from dAIR samples using a commercial enzyme cocktail. The across-environment best linear unbiased estimates (BLUEs) revealed a broad and continuous distribution of saccharification efficiency among the 239 inbred lines (**Figure 1A**). The highest-performing lines ranged from 93.13 to 165.91 mg L⁻¹, exhibiting a remarkable average 4-fold increase in digestibility compared with the lowest-performing entries, which clustered between 22.54 and 28.25 mg L⁻¹. This extensive absolute range (143.37 mg L⁻¹) was accompanied by a 32.25% coefficient of variation (**Table S1**), confirming that biomass recalcitrance is highly divergent within this natural population.

**Figure 1.**
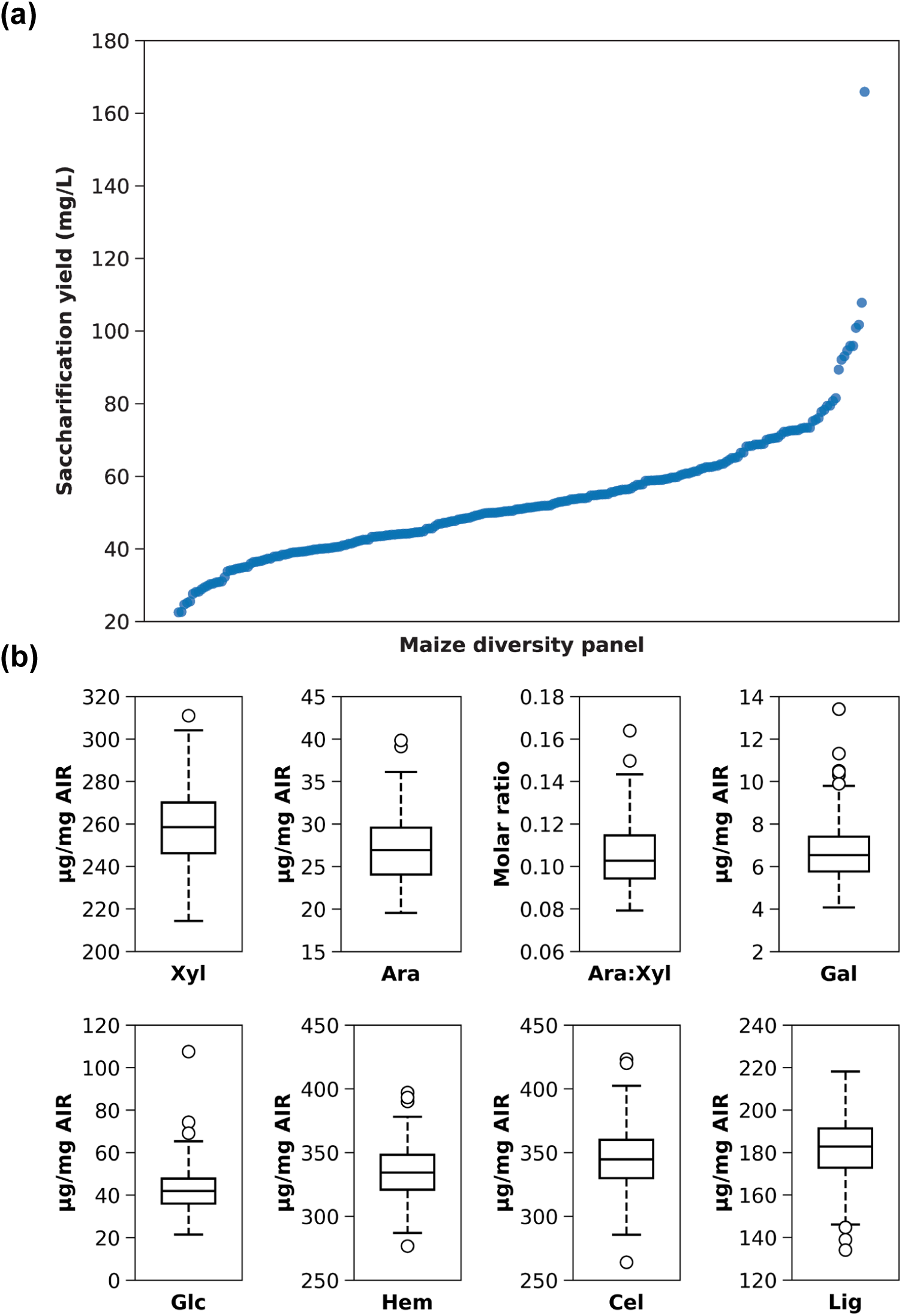
Natural variation in saccharification efficiency and lignocellulosic components among the 239 maize inbred lines. (a) Distribution of saccharification yield. Inbred lines are sorted in ascending order based on the BLUEs of saccharification yield to illustrate the continuous phenotypic variation. (b) Variation in lignocellulosic monosaccharides and polymers. Box plots show the BLUEs of the absolute contents of xylose (Xyl), arabinose (Ara), galactose (Gal), glucose (Glc), hemicellulose (Hem), cellulose (Cel), lignin (Lig), as well as the Ara:Xyl molar ratio. In each box plot, the central line represents the median, the box boundaries indicate the 25th and 75th percentiles, whiskers extend to 1.5× the interquartile range, and open circles denote outliers.

We further characterized the monosaccharide composition of matrix polysaccharides, as well as crystalline cellulose and lignin content of the stalk samples using the across-environment BLUEs for each trait (**Figure 1B and Table S1**). Among the major monosaccharides, xylose (Xyl), the predominant constituent of maize GAX, was the most abundant, ranging from 214.30 to 310.90 mg g⁻¹ cell wall residue. Arabinose (Ara), galactose (Gal), and matrix polymer-derived glucose (Glc) also exhibited considerable variation, ranging from 19.54-39.84, 4.07-13.41, and 21.44-107.50 mg g⁻¹, respectively. In addition, the Ara:Xyl molar ratio, an indicator of GAX substitution and branching, ranged from 0.08 to 0.16. This nearly two-fold variation for this trait suggests broad diversity in the degree of arabinosyl substitution affecting GAX structure. The three major lignocellulosic polymers also displayed diversity within the population. The contents of hemicellulose (Hem, calculated as the sum of constituent monosaccharides), crystalline cellulose (Cel), and lignin (Lig) exhibited significant variations, ranging from 276.70-397.10 mg g⁻¹, 264.00-423.40 mg g⁻¹, and 134.10-218.20 mg g⁻¹, respectively. Notably, crystalline cellulose showed the largest absolute range (159.40 mg g⁻¹) among the structural polymers. Variation was also observed in the relative content of lignocellulosic components (**Figure S1 and Table S1**). Xyl%, Ara%, Gal%, and Glc%, and accounted for 62.27-84.69%, 6.41-11.08%, 1.31-3.88% and 6.85-27.65%, and of the matrix sugars, respectively. Similarly, Hem%, Cel%, and Lig% constituted 32.88-43.22%, 32.31-48.85%, and 15.75-24.80% of total cell wall biomass. Notably, Lig% exhibited higher variation (CV = 7.33%) than either Cel% (CV = 5.79%) or Hem% (CV = 4.86%), indicating greater natural diversity in lignin content which may contribute to the broad variation in cell wall digestibility across the panel.

Taken together, these results revealed substantial chemotypic variation in lignocellulose related traits, indicating extensive genetic diversity for lignocellulosic biomass characteristics across the 239 maize inbred lines and providing a valuable resource for dissecting the genetic architecture of cell wall digestibility and lignocellulosic components.

### Correlation of cell wall digestibility with lignocellulosic components

To investigate how lignocellulosic components contribute to the extensive variation in saccharification efficiency observed at the maize diversity panel level, we examined the trait distribution patterns of both saccharification efficiency and lignocellulosic components, and evaluated their pairwise correlations (**Figure 2 and S2**).

**Figure 2.**
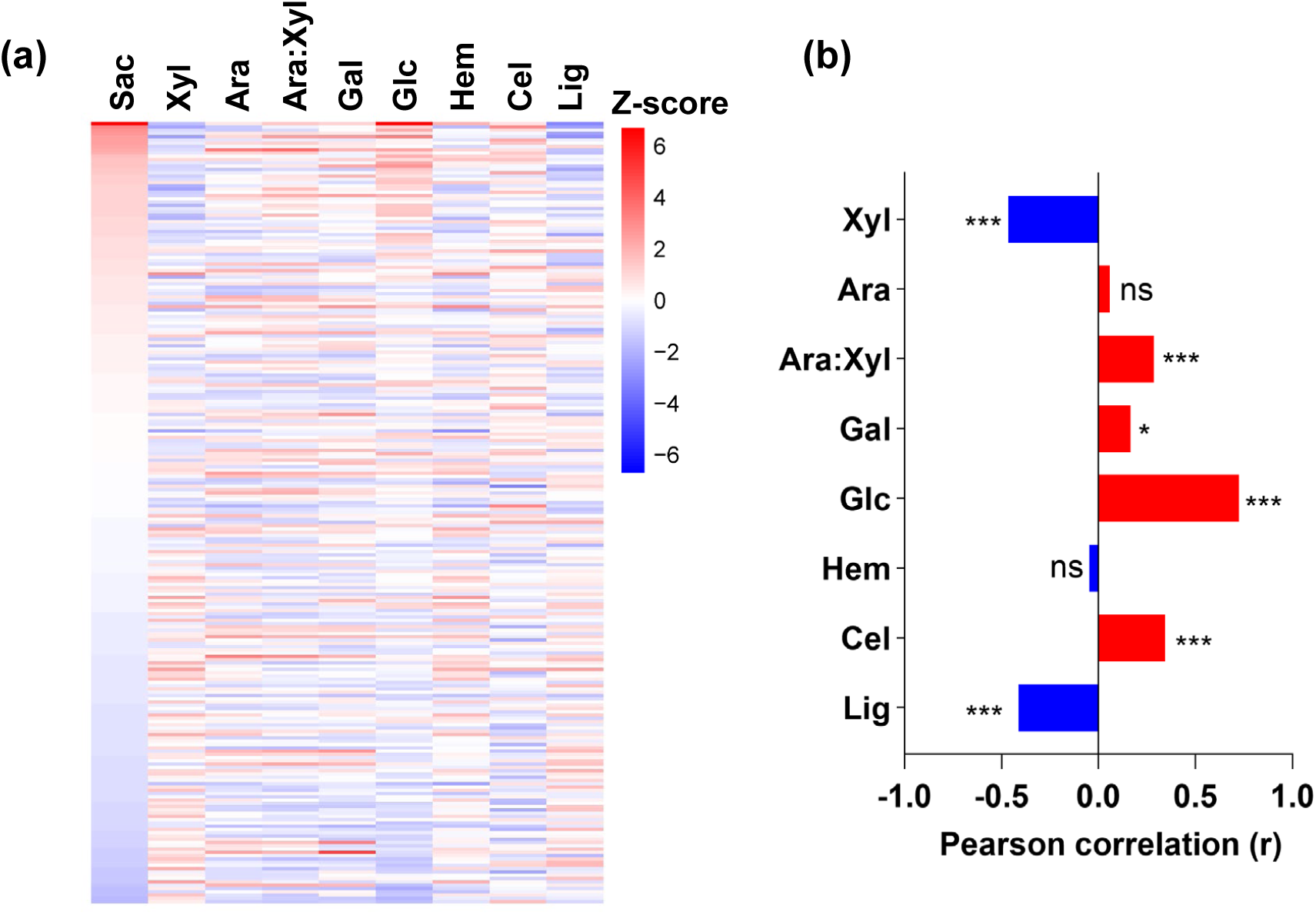
Co-variation and correlation of saccharification efficiency with lignocellulosic components across the maize panel. (a) Z-score heatmap displaying the distribution and co-variation of saccharification efficiency and lignocellulosic components across the maize panel. The 239 inbred lines are ordered by decreasing saccharification efficiency (Sac). BLUEs for each trait were independently standardized to Z-scores. Each row represents one inbred line, and each column represents a trait. The color scale indicates the deviation of each value from the population mean, with red indicating positive Z-scores (values above the population mean) and blue indicating negative Z-scores (values below the population mean). (b) Pearson correlation coefficients (r) between saccharification efficiency and lignocellulosic components. Red and blue bars indicate positive and negative correlations, respectively. Statistical significance is denoted as: ns, no significance; * *p* < 0.05; ** *p* < 0.01; *** *p* < 0.001.

The trait distribution heatmap revealed distinct patterns of co-variation among saccharification efficiency and lignocellulosic components (**Figure 2A and S2A**). Lines exhibiting high saccharification efficiency generally accumulated elevated levels of Glc, Cel, Glc%, Cel%, and Ara:Xyl molar ratio, while tending to display lower levels of Xyl, Lig, Xyl%, and Lig%. Gal and Gal% showed only weak similarity to saccharification patterns, while Ara, Ara%, Hem, and Hem% exhibited no obvious co-distribution with saccharification efficiency.

Correlation analysis further quantified these relationships (**Figure 2B and S2B**). Among all measured traits, matrix glucose content (Glc and Glc%) showed the strongest positive correlation with saccharification efficiency. Similarly, crystalline cellulose content (Cel and Cel%) exhibited a modest positive correlation with digestibility. Among the non-glucose monosaccharides, galactose content (Gal and Gal%) displayed a weak positive correlation with saccharification efficiency, whereas xylose content (Xyl and Xyl%) showed strong negative correlation. However, arabinose (Ara and Ara%) and hemicellulose (Hem and Hem%) contents were not significantly correlated with saccharification efficiency. Interestingly, the Ara:Xyl molar ratio was also positively correlated with saccharification efficiency, suggesting that arabinosyl substitution may also contribute to enhanced biomass digestibility. Consistent with its well-established role as a major contributor to cell wall recalcitrance, lignin content (lignin and lignin%) exhibited a significant negative relationship with saccharification efficiency.

In summary, these results suggest that variation in lignocellulose digestibility at the germplasm level reflects the combined effects of glucose-containing polysaccharides, GAX substitution patterns, and recalcitrance-associated components such as xylan and lignin.

#### Population structure and kinship estimation

To evaluate the population structure and genetic relationships of the 239 maize inbred lines, principal component analysis (PCA) and kinship estimation were performed using genome-wide 44,284 SNP markers (**Figure 3**). The first two principal components explained 8.2% of the genetic variance across the 44,284 SNP markers, and effectively separated the germplasm into three distinct subpopulations (K = 3) (**Figure 3A**). Based on the clustering pattern, 21 lines were assigned to cluster 1, 166 lines to cluster 2, and 52 lines to cluster 3. Cluster 1 positioned on the negative side of PC1, cluster 3 located mainly on the positive side of PC1 and negative side of PC2, and cluster 2 occupying the central region of the PCA plot. The clear separation among clusters indicates the presence of broad genetic diversity and population structure within the panel.

**Figure 3.**
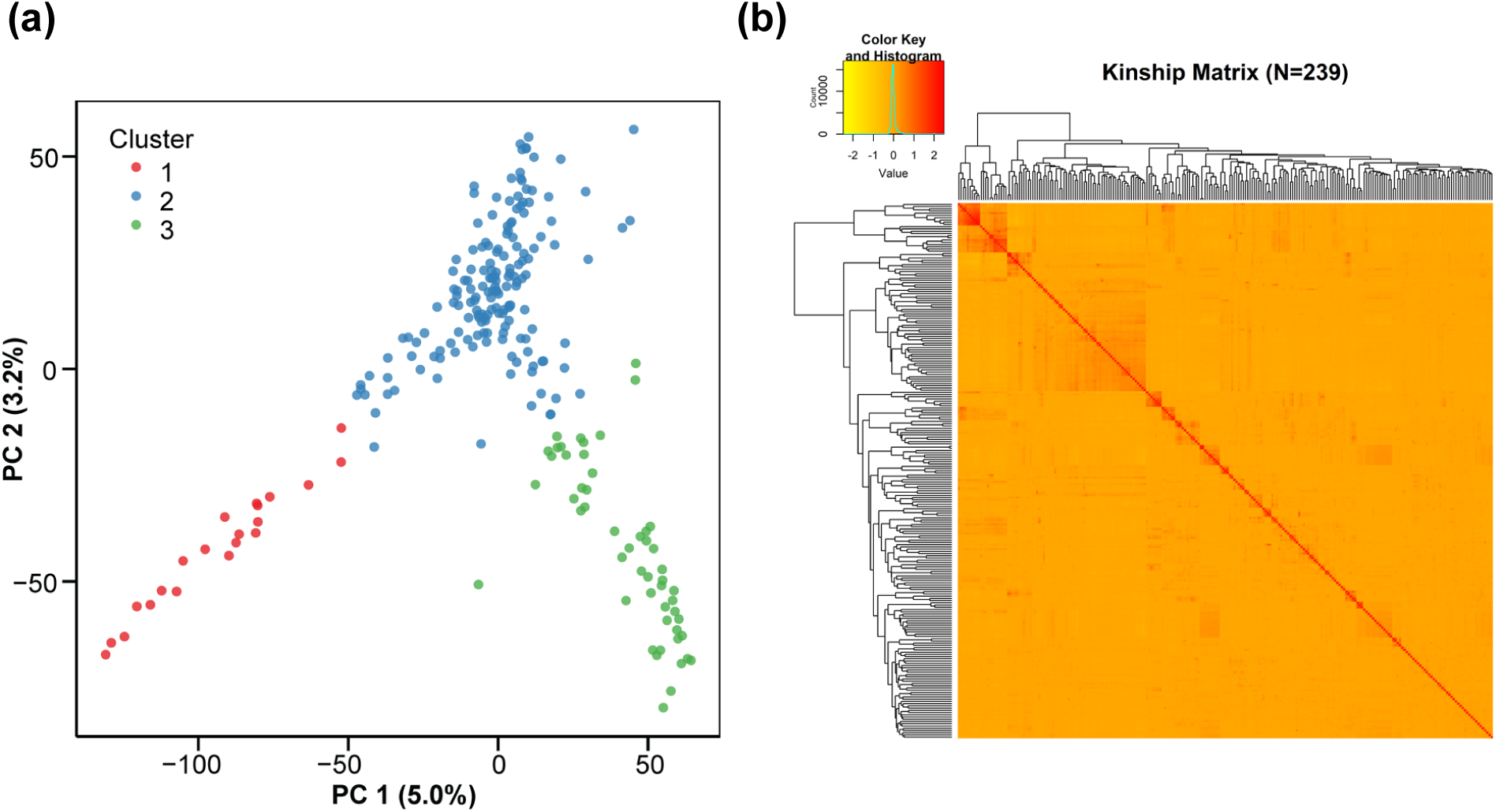
Population structure and kinship analysis of the maize panel. (a) Population structure of the 239 maize inbred lines revealed by principal component analysis (PCA). The x-and y-axis represent the first two principal components (PC1 and PC2), with the percentage of variance explained indicated in parentheses. Lines are colored according to their assigned subpopulations. (b) Heatmap displaying the pairwise kinship matrix among the 239 lines. The color scale ranges from yellow to red, representing low to high kinship coefficients, respectively. Dendrograms on the top and left sides illustrate the hierarchical clustering based on genetic relatedness.

The kinship matrix further characterized the genetic relationships among these inbred lines on a finer level (**Figure 3B**). The heatmap showed that most pairwise kinship coefficients were low, as indicated by the predominance of light orange coloration, suggesting weak relatedness among the majority of accessions. Several small blocks with relatively higher kinship values were observed along the diagonal, reflecting closer genetic relationships among lines within the same subpopulation. However, no extensive regions of strong relatedness were detected, indicating that the panel consisted largely of genetically diverse and unrelated inbred lines.

Overall, the population structure and kinship analyses demonstrated that these 239 inbred lines possess moderate population stratification with three genetic subgroups and generally low levels of familial relatedness. Therefore, both population structure (principal components) and kinship (K matrix) will be incorporated into subsequent GWAS models to effectively control for false-positive associations arising from population stratification and familial relatedness.

### Identification of QTLs associated with saccharification efficiency and lignocellulosic components by GWAS

To identify genetic loci underlying natural variation in digestibility and lignocellulosic component traits, we performed GWAS by fitting a mixed linear model that corrects for population structure and familial relatedness among lines **(Figure 4 and S3)**. Overall, 72 significant SNPs (*P* < 1 x 10^-4^) associated with 16 lignocellulose-related traits were identified across all 10 maize chromosomes, representing 97 significant marker-trait associations (**Table S2**). Among these, 12 SNPs were associated with saccharification efficiency. A total of 55 SNPs were associated with matrix polysaccharide composition, including Ara (4), Ara% (3), Gal (8), Gal% (9), Glc (12), Glc% (8), Xyl (3), and Xyl% (8). In addition, 5 SNPs were associated with the Ara:Xyl molar ratio. Furthermore, 25 SNPs were associated with three lignocellulosic polymers, including hemicellulose (Hem, 3; Hem%, 3), cellulose (Cel, 6; Cel%, 1), and lignin (Lig, 6; Lig%, 6). Notably, 18 of the 72 significant SNPs were associated with two or more traits. For example, SYN4946 was simultaneously associated with Glc, Glc%, Sac, and Xyl% traits.

**Figure 4.**
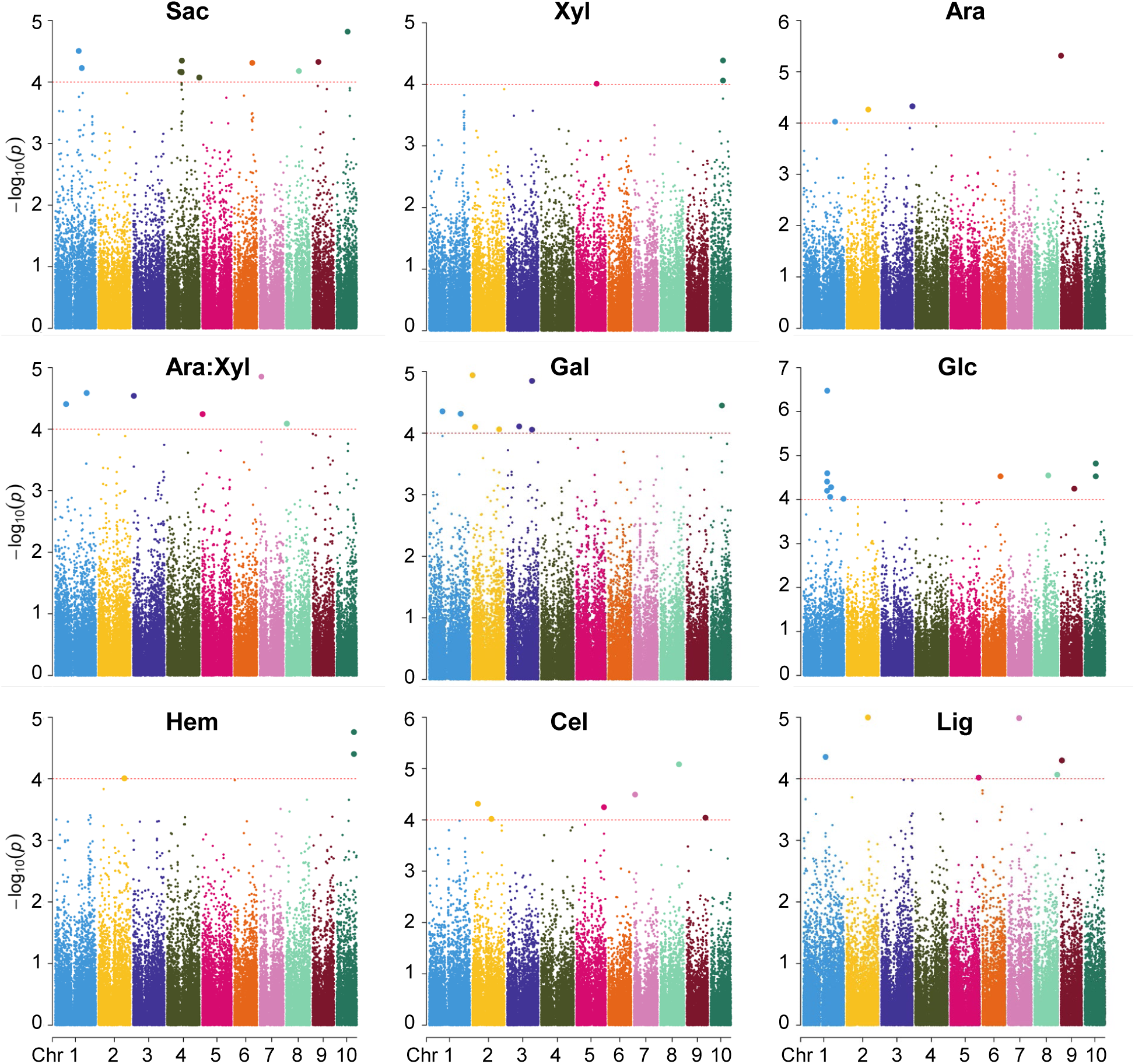
Genome-wide association study identifying significant SNPs associated with saccharification efficiency and lignocellulosic component traits. Points representing SNPs are colored by chromosome. The horizontal red line indicates the suggestive significance threshold -log_10_(*p*) > 4.

Further, a total of 79 QTLs were defined based on integrating the genomic coordinates of significant SNPs, chromosome-specific linkage disequilibrium (LD) decay distances, and pairwise LD among significant SNPs (**Figure 5 and Table S3**). Among them, eight QTLs each were identified for saccharification efficiency and Glc, followed by seven QTLs each for Glc%, Gal and Gal%. Six QTLs each were identified for Xyl%, cellulose, and lignin, whereas five QTLs were detected for Ara:Xyl molar ratio. Four QTLs were identified for Ara and lignin%, three each for Ara% and hemicellulose%, two each for Xyl and hemicellulose, and one for cellulose%.

**Figure 5.**
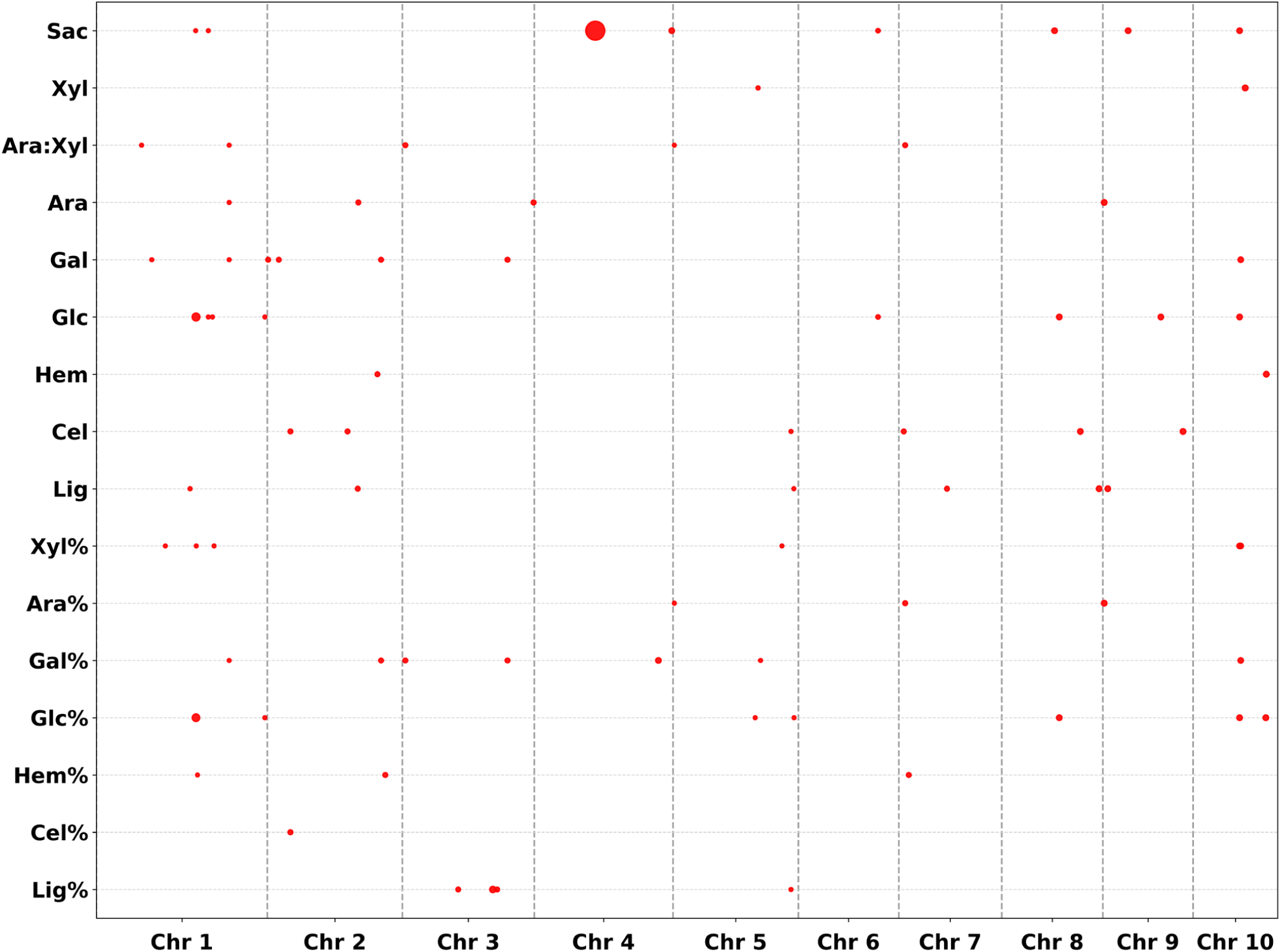
Chromosomal distribution of QTLs associated with saccharification efficiency and lignocellulosic component traits identified in the maize panel. The x-axis indicates the 10 chromosomes, and the y-axis lists the specific traits. Vertical dashed lines delineate boundaries between chromosomes. The area of each circle corresponds to the width of the QTL confidence interval. For QTL nomenclature, the number following the first underscore indicates the chromosome number, and, when present, the final number indicates the sequential index of the QTL on that chromosome. Detailed information regarding the QTLs is provided in Table S3.

The 79 QTLs were unevenly distributed across the 10 maize chromosomes. Chromosome 1 harbored the largest number of QTLs (19), followed by chromosomes 2 (11), 5 (10), 10 (10), and 3 (8), whereas chromosomes 4, 6, 7, 8, and 9 contained relatively fewer loci (2-6 per chromosome). The enrichment of QTLs on chromosomes 1, 2, 3, 5, and 10 suggests that these genomic regions may represent genetic ‘hotspots’ controlling digestibility and lignocellulosic component traits. Furthermore, 43 of the 79 QTLs showed co-localization across at least two different traits. For instance, a major locus on chromosome 10 (designated qSac_10) co-localized with QTLs controlling Glc (qGlc_10), Glc% (qGlc%_10_1), and Xly% (qXyl%_10_1). These cross-trait co-localizations spanning saccharification efficiency, matrix polysaccharides composition, GAX substitution, and lignocellulosic polymer content suggest potential pleiotropic effects or tight genetic linkage among the loci, supporting the presence of shared genetic factors underlying lignocellulosic components and saccharification efficiency.

## Discussion

Lignocellulosic biomass represents an abundant renewable resource for bioenergy production, yet its efficient conversion to fermentable sugars is constrained by the complex and recalcitrant structure of the material (Jørgensen et al., 2007; Marriott et al., 2016). Understanding the genetic determinants of lignocellulosic recalcitrance is therefore important for both elucidating the factors that influence biomass digestibility and improving biomass utilization. GWAS has become a widely used approach for dissecting the genetic basis of lignocellulosic recalcitrance, owing to its capacity to survey naturally occurring genetic variation genome-wide and to identify loci associated with complex quantitative traits at high resolution. Successful applications have been reported across several crop species: in rice, QTLs for saccharification, cellulose, and lignin content have been mapped in diverse rice genotypes (Nguyen et al., 2020; Panahabadi et al., 2022); in sorghum, significant SNPs associated with cellulose, hemicellulose, and lignin content have been identified (Rayaprolu et al., 2021); and in wheat, QTLs for cellulose crystallinity as well as cellulose, hemicellulose, and lignin content have been characterized (Li et al., 2024; Dong et al., 2025). In maize, QTLs for saccharification efficiency, cellulose, hemicellulose, lignin, and cell wall-bound hydroxycinnamate content have been identified across diverse genetic populations, including multi-parent-derived recombinant inbred lines (RILs) and diverse inbred lines (Li et al., 2016; Lopez-Malvar et al., 2019; López-Malvar et al., 2021; Ren et al., 2026). All these GWAS studies demonstrate substantial genetic variation in multiple aspects of lignocellulosic biomass composition and digestibility, and highlight the polygenic nature of biomass recalcitrance.

Despite these advances, several aspects of the genetic basis of maize biomass recalcitrance remain insufficiently explored. First, saccharification efficiency and lignocellulosic component traits were typically evaluated in separate populations or independent experiments, limiting the assessment of their genetic relationships within a unified framework. Second, the previous maize GWAS studies have focused on a limited lignocellulosic-related traits spectrum, such as cellulose, hemicellulose, and lignin content, with less attention to matrix monosaccharide composition and structural features such as the Ara:Xyl molar ratio, which reflects the degree of arabinosyl substitution of xylan. Third, previous maize GWAS studies analyzed a limited set of diversity panels, leaving substantial natural variation in maize germplasm unexplored. To address these gaps, our study simultaneously evaluated saccharification efficiency and a broad range of lignocellulosic component traits in 239 maize inbred lines, most of which had not been previously characterized for these traits **(Figure 1 and S1)**. This integrated design enabled us to examine the genetic relationships between biomass digestibility and multiple aspects of lignocellulosic components within a common genetic and chemotypic framework. Consequently, we identified eight QTLs for saccharification efficiency and 71 QTLs for 15 lignocellulosic component traits **(Figure 5 and Table S3)**. None of the QTLs identified in our study overlapped with those previously reported in maize GWAS. This lack of overlap may arise from several factors, including differences in germplasm diversity, marker density, allele frequency, linkage disequilibrium patterns, chemotyping protocols, and statistical thresholds.

The relationship between lignocellulosic components and saccharification efficiency has been widely investigated across plant species, but the effects of individual lignocellulosic components appear to vary considerably among studies. Consistent with the well-established role of lignin as a major contributor to biomass recalcitrance, we also observed an inverse covariation and strong negative correlation between lignin content and saccharification efficiency in our maize panel **(Figure 2 and S2)**, further highlighting lignin content as a key determinant of natural variation in maize biomass. In contrast, the relationships of cellulose and hemicellulose content with saccharification efficiency appear to be more context dependent. For example, Xu et al. (2012) reported that hemicellulose was the dominant positive factor associated with biomass digestibility, whereas cellulose and lignin showed synergistic negative effects following NaOH or H₂SO₄ pretreatment in Miscanthus samples. Similarly, Li et al. (2015) found that cellulose content was negatively associated with saccharification efficiency, whereas hemicellulose content was positively associated with saccharification following NaOH or H₂SO₄ pretreatment in rice cell wall mutants. In contrast, Pei et al. (2016) found no significant correlation between cellulose content and enzymatic saccharification, whereas hemicellulose content was negatively correlated with saccharification following H₂SO₄- or lime pretreatment in rapeseed straw. In our diverse maize panel, we observed a modest positive correlation between cellulose content and saccharification efficiency, whereas hemicellulose content showed no significant correlation with saccharification **(Figure 2 and S2)**. These contrasting observations suggest that the contribution of cellulose and hemicellulose to biomass digestibility cannot be considered independently of the broader cell wall context and may be shaped by differences in the biomass composition, architecture, and physicochemical properties, as well as by pretreatment and enzymatic hydrolysis conditions.

When examining the relationship between matrix monosaccharide composition and saccharification efficiency **(Figure 2 and S2)**, we found that arabinose (Ara) content showed no significant correlation with saccharification efficiency, whereas galactose (Gal) content showed only a weak positive correlation in our maize panel. These observations also contrast with previous findings in other plant species. Pei et al. (2016) reported positive correlations of both Ara and Gal contents with enzymatic saccharification in H₂SO₄- and lime pretreated rapeseed straw. Similarly, Li et al. (2015) found a positive correlation between Ara content and saccharification efficiency in rice cell wall mutants, attributing this relationship to a negative effect of Ara on cellulose crystallinity. In contrast, Hu et al. (2021) reported a significant negative correlation between Gal content and enzymatic saccharification in rice straw. Such contrasting observations suggest that the contribution of arabinose and galactose to biomass digestibility may depend on differences in species-specific cell wall composition and architecture. Importantly, we observed that matrix-derived glucose content correlated strongly and positively with saccharification efficiency, whereas xylose content showed a strong negative correlation, a pattern that, to our knowledge, has not been previously reported. These findings suggest that variation in glucose-containing matrix polysaccharides and GAX content contribute to maize biomass digestibility. In addition, we also observed a positive correlation between the Ara:Xyl molar ratio and saccharification efficiency in our maize panel, consistent with the similar findings in other plant species (Li et al., 2013; Jia et al., 2014; Li et al., 2025). A higher Ara:Xyl ratio indicates a greater degree of arabinosyl substitution relative to the xylan backbone, which may alter xylan organization and its interactions with other lignocellulosic polymers, potentially enhancing polymers accessibility to hydrolytic enzymes. Together, our findings further advance the understanding of the roles of matrix polysaccharide composition and structural features in shaping biomass recalcitrance.

In conclusion, our study reveals extensive natural variation in saccharification efficiency and lignocellulosic components across a diverse panel of largely undescribed maize germplasm. This variation provides a valuable resource for dissecting the genetic architecture of biomass recalcitrance. Notably, lines exhibiting extreme saccharification efficiency can serve as valuable genetic resources for future breeding aimed at reducing biomass recalcitrance and enhancing biomass utilization. Furthermore, our findings underscore the important roles of glucose-containing polysaccharides, GAX substitution pattern, and recalcitrance-associated components such as xylan and lignin in determining maize biomass digestibility. Finally, the 79 newly identified QTLs provide additional genetic entry points for unraveling the molecular mechanisms underlying lignocellulosic recalcitrance and offer candidate loci for improving maize biomass digestibility.

## MATERIALS AND METHODS

### Maize materials

A total of 239 maize inbred lines were grown at three locations:

1. Cologne-Vogelsang, North Rhine-Westphalia, planting date 21.5.2021, 10 plants per row, 1.5 m row length (2 m including alley). Sampling was performed 6 weeks after flowering.
2. Grissheim, Baden-Württemberg, Planting date 14.5.2021, 10 plants per row, 1.4 m row length (2 m including alley). Sampling occurred 6 weeks after flowering.
3. Einbeck, Southern Lower Saxony, Planting date 10.5.2021, 20 plants per row, 3.2 m row length (4 m including alley). Sampling occurred 6 weeks after flowering.

At maturity, the internode immediately below the first ear was harvested from each inbred line and used for subsequent analyses. The names of the 239 inbred lines and the numbers of biological replicates (individual stems) collected for each line grown across three environments are provided in **Table S4**.

### Cell wall isolation and saccharification assay

Mature stalk samples were prepared as previously described (Wang et al., 2023). Briefly, the dried mature stalks were cut into small pieces and ground into crude powder in a GM200 mixer mill (Retsch). The resulting powder was freeze-dried for 48 h using a Coolsafe system (Scanvac) and subsequently homogenized in 2 mL screw-cap tubes using an MM400 mixer mill (Retsch). De-starched alcohol-insoluble residue (dAIR) was then prepared as previously described (Foster et al., 2010b).

For the saccharification assay, 1 mg of dAIR was incubated with 0.5 µL of Accelerase 1500 (Genencor) in 50 mM citrate buffer (pH 4.5) supplemented with 0.01% NaN₃. A single 5 mm diameter steel ball was added to each sample. Enzymatic hydrolysis was performed for 20 h at 50°C with constant shaking at 250 rpm. The glucose released was quantified using a YSI 2900 Biochemistry Analyzer according to the manufacturer’s instructions.

### Monosaccharide composition and crystalline cellulose content

Monosaccharide composition and crystalline cellulose content were determined simultaneously using a modified one- and two-step sulfuric acid hydrolysis method (Yeats et al., 2016). Briefly, paired dAIR samples (1 mg each) were subjected to either 4% or 72% (w/v) sulfuric acid hydrolysis. For the 4% sulfuric acid hydrolysis, 1.45 mL of 4% (w/v) H₂SO₄ was added directly to one sample. For the 72% sulfuric acid hydrolysis, 50 µL of 72% (w/v) H₂SO₄ was added to the second sample. After incubation at room temperature for 1 h, 1.4 mL of deionized water was added to the 72% hydrolysis sample to reduce the sulfuric acid concentration to 4% (w/v). Both samples were then autoclaved at 121°C for 1 h. Following autoclaving, the hydrolysates were diluted 1:5 and 1:7, respectively, with 0.5 M sodium acetate (pH 5.2) for neutralization.

The resulting monosaccharides were quantified by high-performance anion-exchange chromatography with pulsed amperometric detection (HPAEC-PAD) using a Knauer system. Separation was performed on a CarboPac PA20 analytical column (3 × 150 mm; Thermo Fisher Scientific) equipped with a CarboPac PA20 guard column (3 × 30 mm; Thermo Fisher Scientific). Neutral sugars were separated under isocratic conditions with 2 mM NaOH, followed by a high-strength NaOH wash (700 mM) and re-equilibration with 2 mM NaOH. The analysis was performed at a flow rate of 0.4 mL min⁻¹ and a constant column temperature of 40°C.

Monosaccharides were quantified against external calibration standards of known concentrations, with ribose used as an internal standard. Crystalline cellulose content was calculated by subtracting the glucose concentration derived from 4% sulfuric acid hydrolysis (Glc_matrix) from the total glucose concentration obtained by 72% sulfuric acid hydrolysis hydrolysis (Glc_total). The resulting glucose concentration was multiplied by an anhydrous correction factor of 0.9 to account for the difference in molecular mass between glucose (180.16 g/mol) and anhydroglucose units (162.14 g/mol).

### Total lignin content

Total lignin content was determined using the acetyl bromide soluble lignin (ABSL) assay as previously described (Foster et al., 2010a). Briefly, 1 mg of dAIR was incubated with acetyl bromide reagent (25% [v/v] acetyl bromide in glacial acetic acid) at 50°C for 3 h. Following incubation, 2 M NaOH and 0.5 M hydroxylamine hydrochloride were added, and the final volume was adjusted to 2 mL. Absorbance was measured at 280 nm against a reagent blank using a spectrophotometer. The percentage of ABSL was calculated according to the following equation:

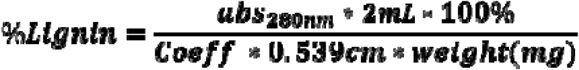

where Coeff is the grass-specific empirical coefficient (17.75), and 0.539 cm is the optical path length of the 96-well plate.

### Best linear unbiased estimates

To account for environmental and experimental variation, a best linear unbiased estimate (BLUE) of each trait across all environmental trials was obtained for each inbred line using the *lme4* package (Bates et al., 2015) in R (version 4.2.2). A linear mixed model was fitted with line as a fixed effect and environment, line × environment interaction, and plot as random effects:

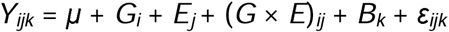

where *Y_ijk_* represents the observed phenotypic value, μ is the overall mean, *G_i_* is the effect of the *i*th inbred line, *E_j_* is the effect of the *j*th environment, (*G* × *E*)*_ij_* is the interaction between the *i*th inbred line and the *j*th environment, *B_k_* is the bock effect, and ε*ijk* is the residual error. Adjusted means for each inbred line were estimated using the *emmeans* package (Lenth, 2023) and subsequently used as BLUEs for statistical analyses, correlation analysis, and genome-wide association studies (GWAS).

### Genome-wide association study

The 239 maize inbred lines were genotyped with the Illumina MaizeSNP50 BeadChip containing 56,110 SNPs. After quality control, a total of 44,284 high-quality SNPs with a minor allele frequency (MAF) ≥0.05 were retained for subsequent analyses. Based on these SNPs, principal component analysis (PCA) and calculation of pairwise kinship matrix (K matrix) were performed. The BLUEs for inbred lines were used as phenotypic inputs for GWAS. GWAS was performed using a mixed linear model (MLM) implemented in the *sommer* package (Covarrubias-Pazaran, 2016) in R (version 4.2.2). In the model, the first two principal components (PC1 and PC2) and the K matrix were included as fixed and random effects, respectively, to account for population structure and genetic relatedness. Significant associations were detected at a suggestive threshold of −log_10_(*P*) > 4.0 (*P* < 1.0 × 10^−4^) (Li et al., 2016; Guo et al., 2023).

### Linkage disequilibrium analysis, QTL definition, and candidate gene identification

The 44,284 high-quality SNPs were assigned to their respective chromosomes, and chromosome-specific linkage disequilibrium (LD) was estimated separately for each chromosome using TASSEL 5.0 software (Bradbury et al., 2007). LD decay plots were generated by plotting pairwise *r^2^* values against the physical distance between SNPs in R (version 4.2.2) **(Figure S4)**.

For each significant SNP identified by GWAS, the genomic coordinates were converted from the B73 RefGen_v2 assembly to the B73 RefGen_v5 assembly (Matthew et al., 2021). QTL intervals were then defined by extending the genomic region upstream and downstream of each significant SNP according to the chromosome-specific LD decay distance at *r^2^* = 0.1. Overlapping QTL intervals were merged and considered a single QTL. Genes located within the resulting QTL intervals were extracted from the Ensembl Plants database and subsequently annotated at MaizeMine v1.6 database.

## AUTHOR CONTRIBUTIONS

VR and MP conceived and designed the project. BS, DVI, FR, SG and TB designed the field trial and collected stalk sample. BS wrote R scripts and provided SNP array data for GWAS analysis. SW performed experiments and analyzed data. SW, MP and VR wrote the manuscript. All authors contributed to the article and approved the submitted version.

## ACKNOWLEDGMENTS

The authors would like to thank Katharina Grosche and Barbara Schulten for excellent technical support in cell wall analyses. This work was funded by Germany’s Federal Ministry of Education and Research (BMBF) grant “Cornwall”, 031B1303A.

## CONFLICT OF INTEREST

Susanne Groh and Frank Röber are both employed by Pioneer Hi-Bred. Therese Bolduan is employed by KWS-Saat.

## DATA AVAILABILITY STATEMENT

All data supporting the findings of this study are available within the paper and its Supporting Information.

## SUPPORTING INFORMATION

**Figure S1.** Distribution of relative abundance of lignocellulosic monosaccharides and polymers among the 239 maize inbred lines. Box plots show the BLUEs of the relative abundance of xylose (Xyl%), arabinose (Ara%), galactose (Gal%), glucose (Glc%), hemicellulose (Hem%), cellulose (Cel%), and lignin (Lig%). In each box plot, the central line represents the median, the box boundaries indicate the 25th and 75th percentiles, whiskers extend to 1.5× the interquartile range, and open circles denote outliers.

**Figure S2**. Co-variation and correlation analysis of saccharification efficiency and relative abundances of lignocellulosic monosaccharides and polymers.

(a) Z-score heatmap showing the distribution and co-variation of saccharification efficiency and relative abundances of lignocellulosic monosaccharides and polymers. The 239 inbred lines are ordered by decreasing saccharification efficiency (Sac). BLUEs for each trait were independently standardized to Z-scores. Red indicates positive Z-scores, whereas blue indicates negative Z-scores.

(b) Pearson correlation coefficients (r) between saccharification efficiency and relative abundances of lignocellulosic monosaccharides and polymers. Red and blue bars indicate positive and negative correlations, respectively. Statistical significance is denoted as: ns, no significance; * *p* < 0.05; ** *p* < 0.01; *** *p* < 0.001.

**Figure S3**. Manhattan plots showing significant associations with the relative abundance of lignocellulosic monosaccharides and polymers. Points representing SNPs are colored by chromosome. The horizontal red line indicates the suggestive significance threshold -log10(*p*) > 4.

**Figure S4**. Linkage disequilibrium (LD) decay distance across the 10 maize chromosomes. Each panel represents a specific chromosome, with the red curve indicating the fitted LD decay model, and the blue dashed line marking the physical distance where LD decays to *r^2^* = 0.1. The specific LD decay distance (in kb) for each chromosome is annotated at the top of each panel. The Y-axis represents the squared correlation coefficient (*r^2^*) as a measure of pairwise LD, and the X-axis indicates the physical distance in base pairs (kb).

**Table S1**. Phenotypic variations for saccharification efficiency and lignocellulosic component traits among the 239 maize inbreds.

**Table S2**. List of 97 significant SNP-trait associations detected by GWAS.

**Table S3**. List of 79 QTLs associated with saccharification efficiency and lignocellulosic component traits.

**Table S4**. List of the 239 inbred lines and the numbers of biological replicates collected for each line.

## Notes

### Competing Interest Statement

The authors have declared no competing interest.

