## Supplemental Figures 1-3 for "Genome-wide association study in a maize diversity panel dissects the genetic basis of lignocellulosic recalcitrance"

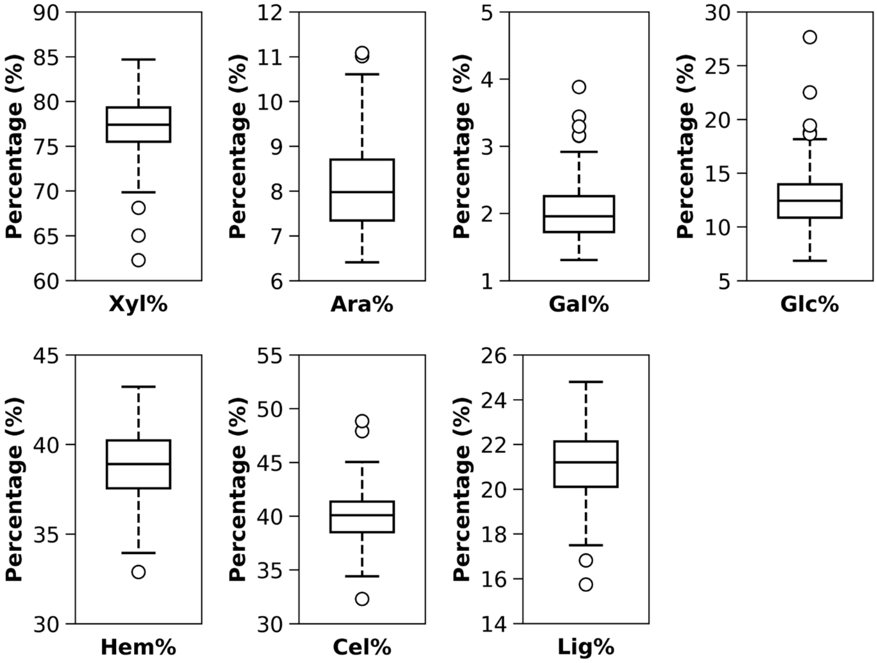


**Supplementary Figure 1**. Distribution of relative abundance of lignocellulosic monosaccharides and polymers among the 239 maize inbred lines. Box plots show the BLUEs of the relative abundance of xylose (Xyl%), arabinose (Ara%), galactose (Gal%), glucose (Glc%), hemicellulose (Hem%), cellulose (Cel%), and lignin (Lig%). In each box plot, the central line represents the median, the box boundaries indicate the 25th and 75th percentiles, whiskers extend to 1.5× the interquartile range, and open circles denote outliers.

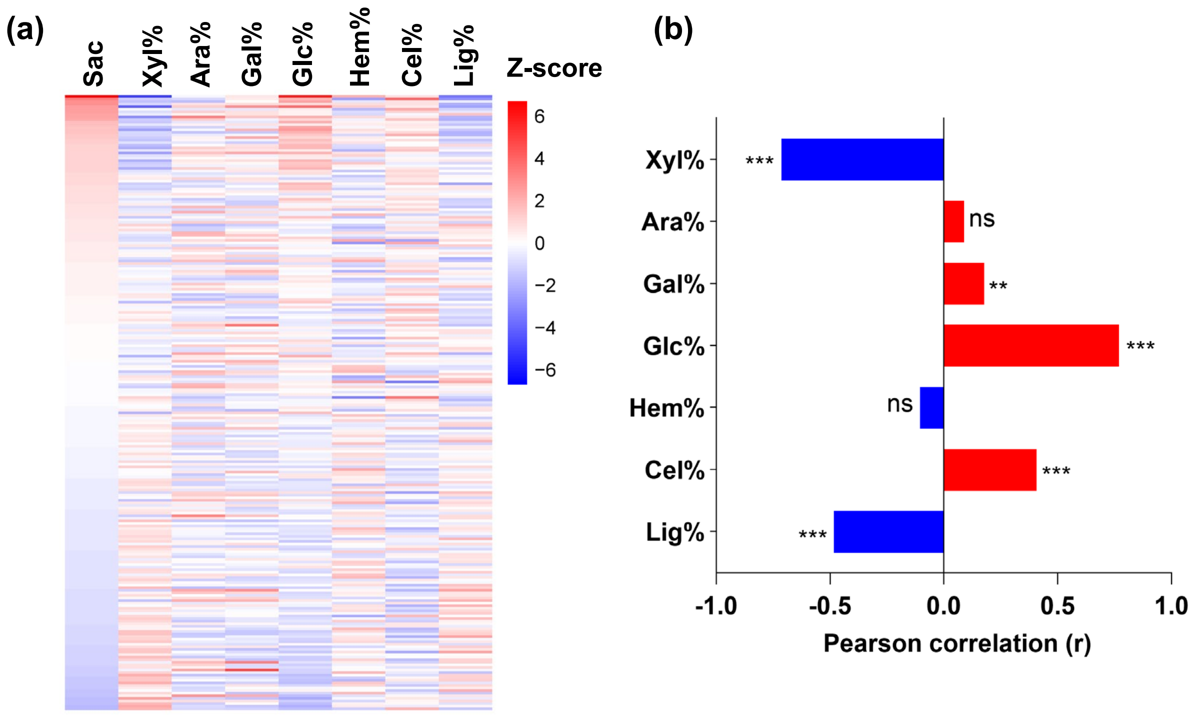


**Supplementary Figure 2**. Co-variation and correlation analysis of saccharification efficiency and relative abundances of lignocellulosic monosaccharides and polymers.

(a) Z-score heatmap showing the distribution and co-variation of saccharification efficiency and relative abundances of lignocellulosic monosaccharides and polymers. The 239 inbred lines are ordered by decreasing saccharification efficiency (Sac). BLUEs for each trait were independently standardized to Z-scores. Red indicates positive Z-scores, whereas blue indicates negative Z-scores.

(b) Pearson correlation coefficients (r) between saccharification efficiency and relative abundances of lignocellulosic monosaccharides and polymers. Red and blue bars indicate positive and negative correlations, respectively. Statistical significance is denoted as: ns, no significance; * $p < 0.05$ ; ** $p < 0.01$; *** $p < 0.001$.

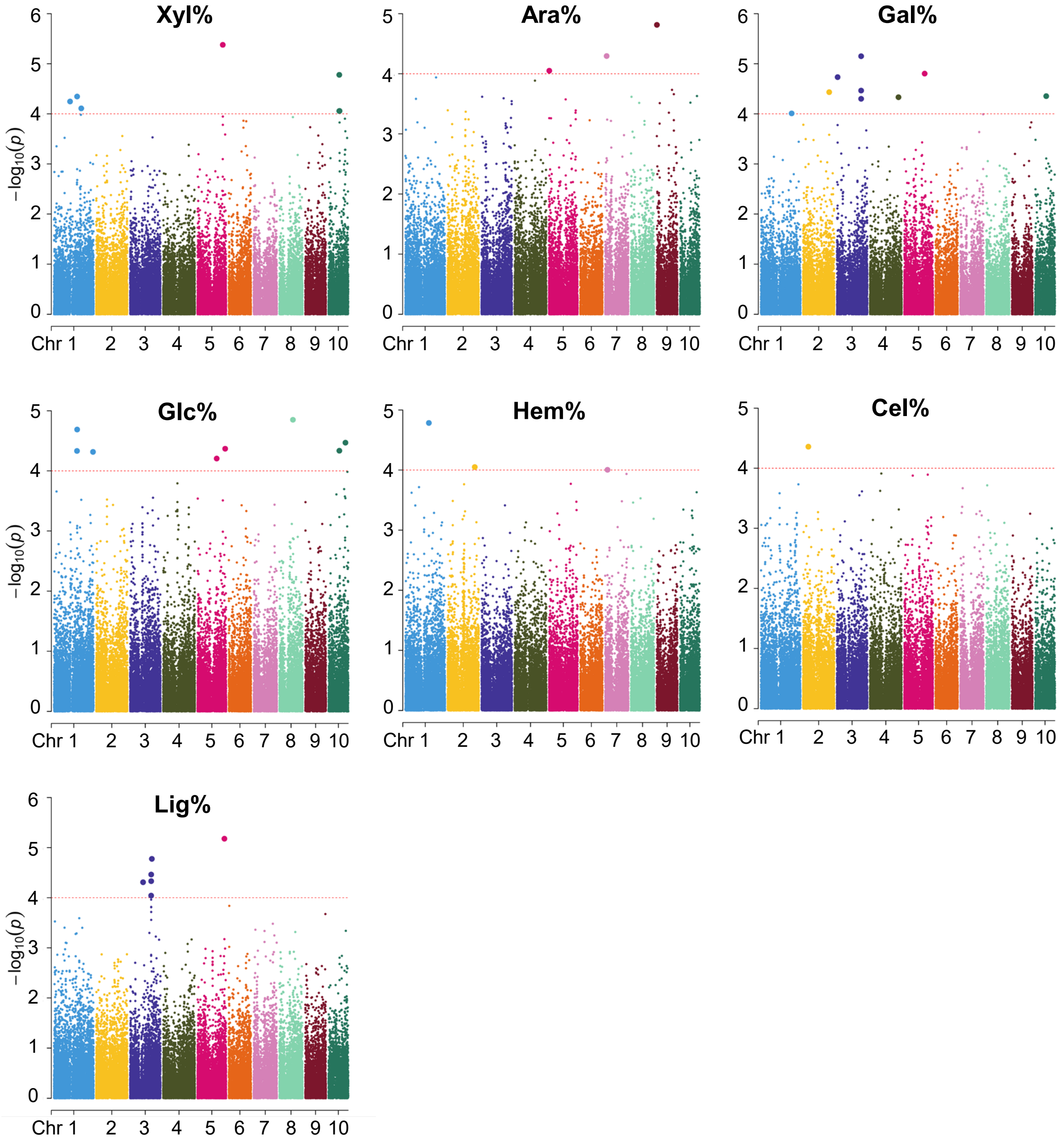


**Supplementary Figure 3.** Manhattan plots showing significant associations with the relative abundance of lignocellulosic monosaccharides and polymers. Points representing SNPs are colored by chromosome. The horizontal red line indicates the suggestive significance threshold $-\log_{10}(p) > 4$.

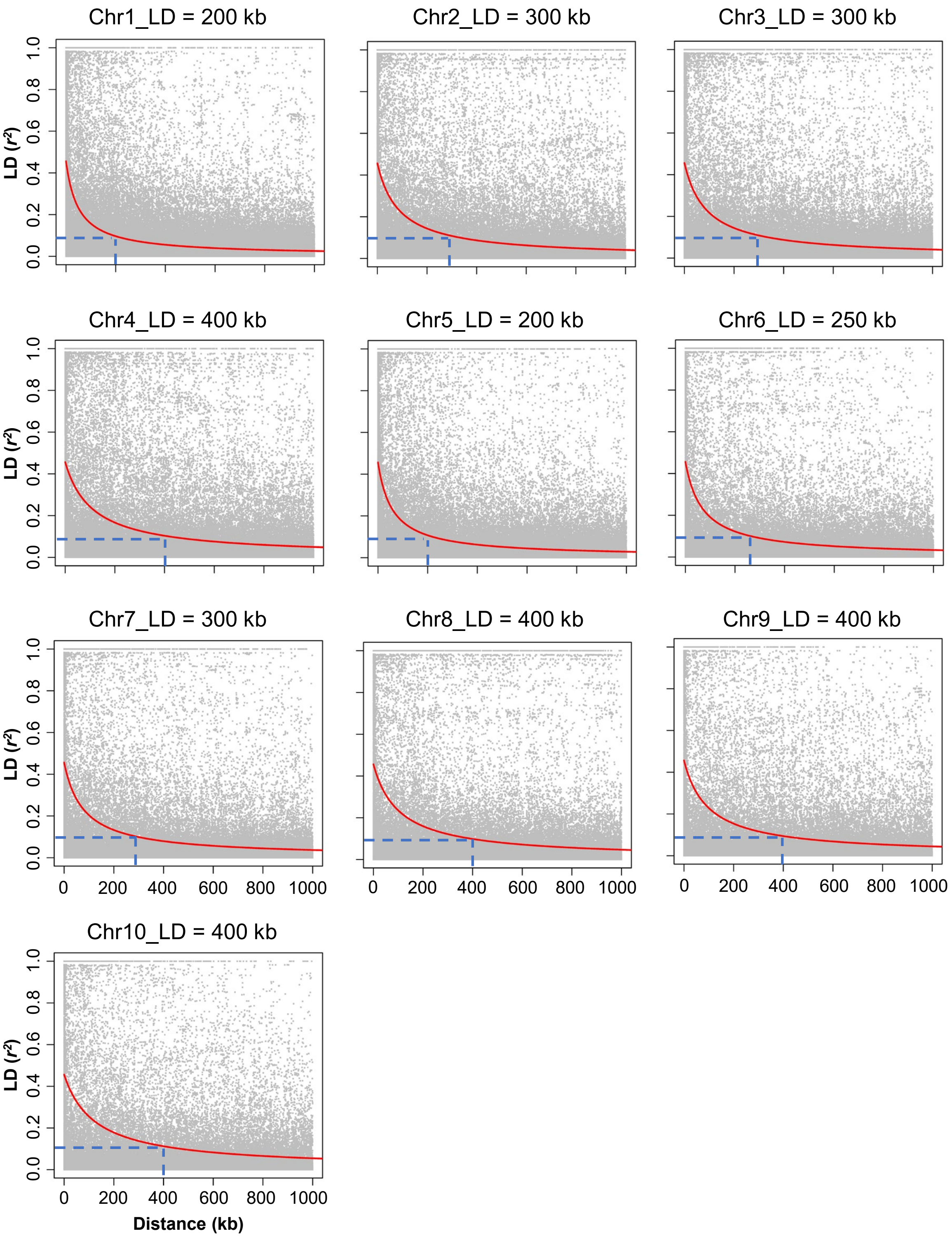


**Supplementary Figure 4**. Linkage disequilibrium (LD) decay distance across the 10 maize chromosomes. Each panel represents a specific chromosome, with the red curve indicating the fitted LD decay model, and the blue dashed line marking the physical distance where LD decays to $r^2 = 0.1$. The specific LD decay distance (in kb) for each chromosome is annotated at the top of each panel. The Y-axis represents the squared correlation coefficient ($r^2$) as a measure of pairwise LD, and the X-axis indicates the physical distance in base pairs (kb).
